# Visual feature tuning of superior colliculus neural reafferent responses after fixational microsaccades

**DOI:** 10.1101/2019.12.24.888149

**Authors:** Fatemeh Khademi, Chih-Yang Chen, Ziad M. Hafed

## Abstract

The primate superior colliculus (SC) is causally involved in microsaccade generation. Moreover, visually-responsive SC neurons across this structure’s topographic map, even at peripheral eccentricities much larger than the tiny microsaccade amplitudes, exhibit significant modulations of evoked response sensitivity when stimuli appear peri-microsaccadically. However, during natural viewing, visual stimuli are normally stably present in the environment and are only shifted on the retina by eye movements. Here we investigated this scenario for the case of microsaccades, asking whether and how SC neurons respond to microsaccade-induced image jitter. We recorded neural activity from two male rhesus macaque monkeys. Within the response field (RF) of a neuron, there was a stable stimulus consisting of a grating of one of three possible spatial frequencies. The grating was stable on the display, but microsaccades periodically jittered the retinotopic RF location over it. We observed clear short-latency visual reafferent responses after microsaccades. These responses were weaker, but earlier (relative to new fixation onset after microsaccade end), than responses to sudden stimulus onsets without microsaccades. The reafferent responses clearly depended on microsaccade amplitude, as well as microsaccade direction relative to grating orientation. Our results indicate that one way for microsaccades to influence vision is through modulating how the spatio-temporal landscape of SC visual neural activity represents stable stimuli in the environment. Such representation strongly depends on the specific pattern of temporal luminance modulations expected from the relative relationship between eye movement vector (size and direction), on the one hand, and spatial visual pattern layout, on the other.

**Significance statement:** Despite being diminutive, microsaccades still jitter retinal images. We investigated how such jitter affects superior colliculus (SC) activity. We found that SC neurons exhibit short-latency visual reafferent bursts after microsaccades. These bursts reflect not only the spatial luminance profiles of visual patterns, but also how such profiles are shifted by eye movement size and direction. These results indicate that the SC continuously represents visual patterns, even as they are jittered by the smallest possible saccades.

## Introduction

Microsaccades (reviewed in Ahissar et al. 2016; Engbert 2006; Hafed 2011; Hafed et al. 2015; Krauzlis et al. 2017; Martinez-Conde et al. 2004; Poletti and Rucci 2016; Rolfs 2009) are small saccades that occur periodically during prolonged gaze fixation. In comparison to larger saccades, with peak velocities as high as ∼800 deg/s, microsaccades (having peak velocities well below 100 deg/s) decidedly cause milder disruptions of retinal images. Nonetheless, these fixational eye movements still jitter images, and they are therefore expected to influence visual neural activity due to the sequence of motion blur followed by stable visual stimulus that they cause. Indeed, microsaccades are associated with post-movement visual reafferent responses in a variety of brain areas (Bosman et al. 2009; Herrington et al. 2009; Kagan et al. 2008; Leopold and Logothetis 1998; Martinez-Conde et al. 2002; 2000; Snodderly et al. 2001). However, the bulk of past work has focused primarily on the mere presence of reafferent responses, but not necessarily on their properties. While such presence may be relevant for issues like the refreshing of retinal images by microsaccades, perhaps to help in counteracting visual fading (Martinez-Conde et al. 2004), understanding the properties of post-microsaccadic visual reafference is additionally important for clarifying the mechanisms of visual neural coding when the sensor for vision (the eye) is continually mobile (Kagan et al. 2008; Kuang et al. 2012; Rucci et al. 2007; Segal et al. 2015; Snodderly et al. 2001).

In this study, we explored the properties of visual reafference after microsaccades in superior colliculus (SC) visual activity. The SC has historically been studied in primate animal models primarily from a saccade motor control perspective (Basso and May 2017; Gandhi and Katnani 2011; Krauzlis et al. 2017; Robinson 1972). Indeed, the SC is causally involved in microsaccade generation (Hafed et al. 2009; Hafed and Krauzlis 2012; Hafed et al. 2013; Willeke et al. 2019). However, the SC, even in primates, also exhibits significant visual responses (Goldberg and Wurtz 1972), and recent results have revealed visual pattern analysis capabilities of SC neurons (Chen and Hafed 2018; Chen et al. 2019; Chen et al. 2018; Hafed 2018; Hafed and Chen 2016; Hall and Colby 2016; Herman and Krauzlis 2017). This leaves the question of how the SC represents continuously presented stimuli (as opposed to stimulus onsets) open, and how such representation is modulated by microsaccades.

In all of our previous investigations of SC neural modulations by microsaccades, we presented stimuli suddenly around the time of these eye movements (Bellet et al. 2017; Chen and Hafed 2017; Chen et al. 2019; Chen et al. 2015; Hafed et al. 2015; Hafed and Krauzlis 2010). Whether and how SC neurons would react to stable visual stimuli being jittered on the retina by microsaccades are questions that have not yet been explored. Investigating these questions would be important to support the notion that the SC may be viewed as much as an early visual area as it is viewed as a late motor control area, and it would also be in line with a potential role for the SC in signaling visual salience and priority in the environment (Veale et al. 2017; White et al. 2017a; White et al. 2019; White et al. 2017b).

In our experiments, we exploited our recent observations of visual feature tuning of SC neurons for different spatial frequencies (Chen et al. 2018), and we compared such tuning to the case when the stimuli were stably present in the environment and only jittered retinotopically by microsaccades. We analyzed the relative relationships between microsaccade sizes and directions, on the one hand, and grating spatial frequencies, on the other; we found that the specific pattern of temporal luminance modulations caused within a given RF by a given microsaccade dictates how the neuron “updates” its post-microsaccadic representation of the otherwise stable stimulus. This suggests that the SC can contribute to visual coding of scenes during active visual fixation.

## Materials and Methods

### Ethics approvals

All monkey experiments were approved by ethics committees at the Regierungspräsidium Tübingen. The experiments were in line with the European Union directives, and the German laws, governing animal research.

### Laboratory setups

Monkey neurophysiology experiments were performed in the same laboratory environment as that described in (Chen et al. 2019; Chen et al. 2015; Chen et al. 2018).

### Animal preparation

We collected behavioral and neural data from 2 adult, male rhesus macaques (macaca mulatta). Monkeys N and P (aged 7 years and weighing 8 and 7 kg, respectively) were implanted with scleral search coils to allow measuring eye movements using the electromagnetic induction technique (Fuchs and Robinson 1966; Judge et al. 1980). The monkeys were also implanted with a head holder to stabilize their head during the experiments, as well as recording chambers to access the right and left SC’s. Details on all implant surgeries were provided earlier (Chen and Hafed 2013; Chen et al. 2015).

### Monkey behavioral tasks

Each monkey performed a simple fixation task. Prior to running this main task (which we describe below), we first assessed the visual response field (RF) of a recorded SC neuron using standard fixation and saccade behavioral paradigms described earlier (Chen and Hafed 2018; Chen et al. 2019; Chen et al. 2015; Chen et al. 2018). Briefly, the monkey sat in a dark room in front of a gray display with luminance 21 cd/m^2^. A white (72 cd/m^2^) spot (square of 8.5 x 8.5 min arc) was then presented at different display locations while the monkey was maintaining gaze fixation on a similar spot. Visual responses to the appearing spot were used to assess the visual RF location and boundaries. In some variants, the monkey also made a later saccade to the stimulus.

We then ran the main task of this study. A similar fixation spot to that described above was first presented at the center of the display for the monkey to fixate. After the monkey fixated the spot for ∼490 ms (+/- 80 ms s.d.), a high-contrast (100%) Gabor grating appeared at a location centered on the recorded neuron’s RF. The radius of the grating was tailored to the size of the RF, and the spatial frequency could be 0.56, 2.22, or 4.44 cycles per degree (cpd). These spatial frequencies are known to be effective in driving visual responses in the SC (e.g. Chen et al. 2018). The grating persisted for ∼1300-3000 ms until trial end. If the monkey maintained fixation until trial end, the monkey was rewarded with a juice/water drop.

In all experiments on both monkeys, we used a vertical Gabor grating. While the rhesus macaque SC does exhibit orientation tuning (Chen and Hafed 2018), our experience has been that tuning curves are broad enough to include notable responses for vertical gratings as well. We thus used vertical gratings in a variety of our recent studies (Chen and Hafed 2017; Chen et al. 2015; Chen et al. 2018). Because microsaccades vary in direction when they occur (e.g. see Fig. 1c in Results), this still allowed us to explore the relative relationships between movement vector direction and stimulus pattern orientation (see Results). We also confirmed this idea further by additionally running one of the monkeys (monkey N) on a variant of the task using horizontal, rather than vertical, gratings. For a majority of neurons recorded from this monkey in this condition (16/28), we ran one block of trials with vertical gratings and another block of trials with horizontal gratings. That is, the same neuron was studied with the two different stimulus orientations.

**Figure 1.**
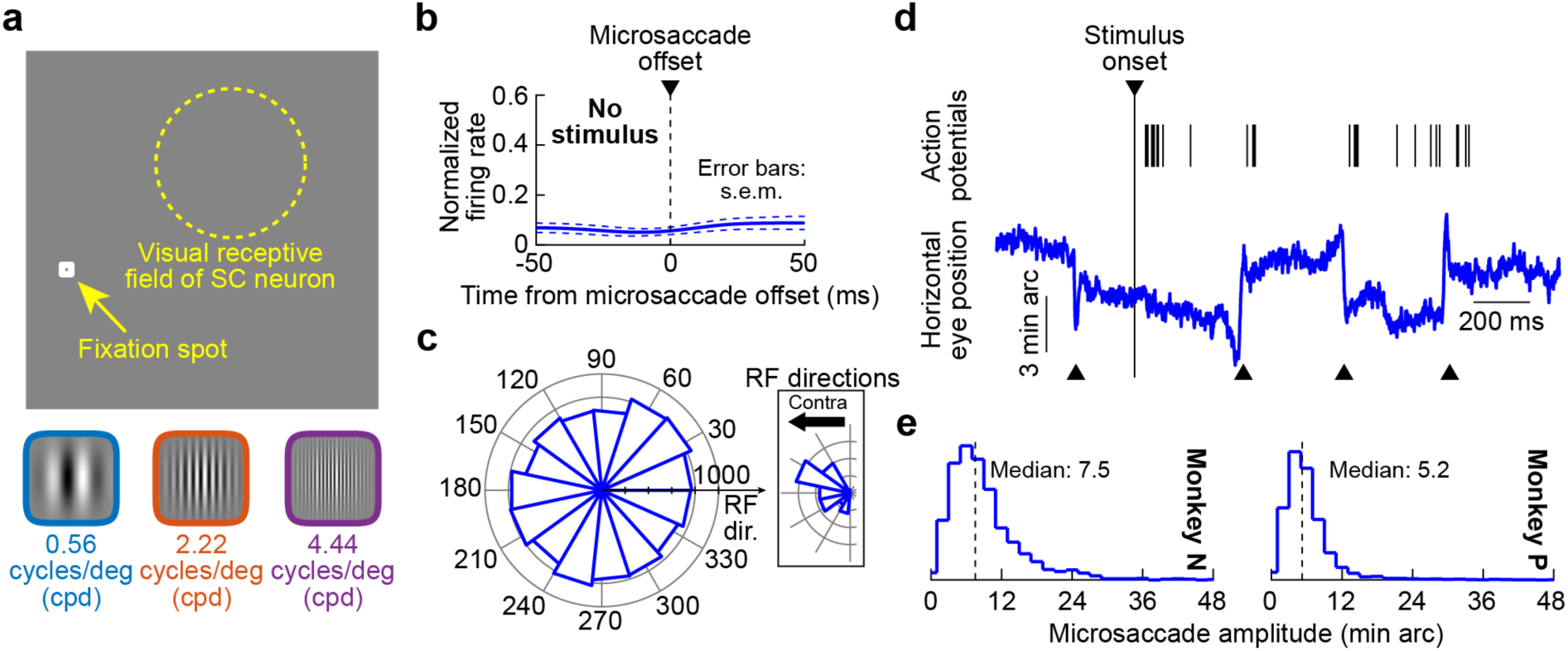
Task design. **(a)** The monkeys fixated, and we presented an RF stimulus comprising of Gabor gratings having different spatial frequencies (schematized below the top panel). **(b)** Without an RF stimulus, our neurons did not exhibit microsaccade-related discharge. We normalized each neuron’s firing rate based on its peak visual evoked response after stimulus onset (e.g. Fig. 5; Materials and Methods), and we then aligned all neurons’ (n=55) normalized activity to microsaccade onset when the microsaccades happened without any RF stimulus present (i.e. during initial fixation). There was no modulation of activity caused by microsaccades in the absence of visual stimulation. **(c)** Microsaccade directions were uniformly distributed relative to RF location across our population (and, therefore, grating orientation since such orientation was fixed across RF’s). This allowed us to investigate different relationships between movement directions and stimulus patterns in our analyses (e.g. Fig. 9) independently of absolute microsaccade direction. The inset shows the range of RF directions in our population. **(d)** Example eye position trace (blue) and neural spiking activity (each black tick mark represents an action potential) from a sample recorded SC neuron. Shortly after stimulus onset, there was a burst of action potentials (the visual evoked response). Subsequently, the neuron occasionally emitted activity, with bursts after microsaccades (visual reafferent responses); the black upward triangles indicate microsaccade onsets. We analyzed such bursts during a sustained fixation interval with the stimulus stably present over the RF (Materials and Methods). **(e)** In both monkeys, microsaccade amplitudes that we analyzed were primarily smaller than 0.2 deg (12 min arc), as shown by the histograms of amplitude probabilities.

**Figure 2.**
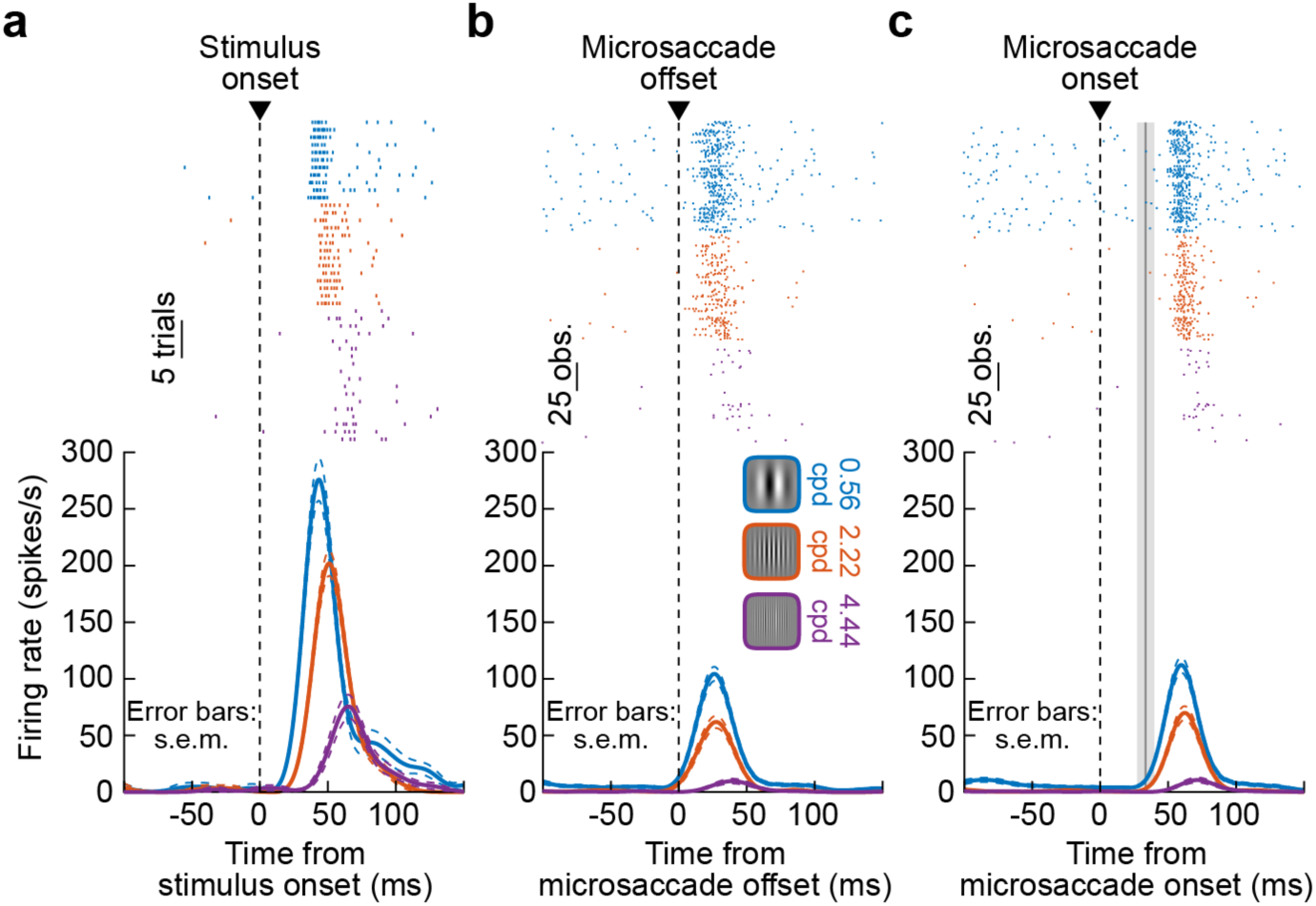
Activity of an example SC neuron after stimulus onset and after microsaccades. **(a)** Spiking activity as a function of time after stimulus onset. Each tick mark in the top panel represents an action potential, and each row of action potentials represents a trial. Trials were sorted by the spatial frequency of the RF stimulus (vertical grating). The bottom panel shows the same data but now plotted as firing rate curves. Error bars denote s.e.m. across trials. The neuron had a stronger and earlier visual burst for the lowest spatial frequency. The numbers of trials per spatial frequency are shown in the top panel. **(b)** For each microsaccade occurring while the grating was visible (that is, in a sustained interval after grating onset; Materials and Methods), we aligned the same neuron’s activity to microsaccade end (labeled “offset” in the figure). The neuron emitted a visual reafferent burst, which also depended on spatial frequency. Note how the visual reafferent response (relative to new fixation onset after a microsaccade) occurred earlier than the stimulus-evoked response in **a**. Error bars in this case denote s.e.m. across the number of microsaccades analyzed. **(c)** Same as in **b** but now aligned to microsaccade onset. The vertical gray line and shaded surrounding interval denote mean microsaccade duration and bounds, respectively.

### Behavioral analyses

We detected saccades and microsaccades using established methods in our laboratory (Bellet et al. 2019; Chen and Hafed 2013). We manually inspected each trial to correct for false alarms or misses by the automatic algorithms, which were rare. We also marked blinks or noise artifacts for later removal. We characterized microsaccade times in order to align neural activity to them (for analyzing visual reafferent responses). We also characterized microsaccade amplitudes (i.e. sizes) and directions in order to explore interactions between movement vector and stimulus spatial frequency.

### Neural firing rate analyses

We analyzed data from 55 well-isolated single neurons (23 from monkey N and 32 from monkey P). These neurons were collected with the vertical gratings. With horizontal gratings, we analyzed 28 neurons from monkey N. Our criteria for analyzing neurons were the presence of a visual evoked response for at least one of the spatial frequencies that we presented, as well as the availability of >10 trials per presented spatial frequency in a given recording. Our numbers of neurons are typical of recent SC studies exploring visual responses (e.g. White et al. 2019; White et al. 2017b), and the response profiles that we observed were sufficiently homogeneous, and consistent with our other recorded tasks in the SC (Chen and Hafed 2017; Chen et al. 2018; Hafed and Chen 2016), to make our sample of neurons representative of SC neurons in general.

The neurons described in this study included both purely visual and visual-motor neurons, but we did not notice substantial differences in the present analyses (focused on visual responses only) between the two types. We therefore did not separate analyses for these two functional types of SC neurons. We did, however, ensure that none of our neurons exhibited movement-related discharge for microsaccade generation (Hafed et al. 2009; Hafed and Krauzlis 2012; Willeke et al. 2019). Such discharge would come at the time of potential visual reafferent responses and mask such responses. Given that our recorded neurons (see Results) were not extremely foveal (Chen et al. 2019), exclusion of microsaccade-related discharge was trivially easy. We also checked this explicitly in Fig. 1b in Results.

We used firing rates from the RF mapping tasks alluded to above in order to confirm that our placement of gratings during the experiments was valid. We then analyzed the neural data in the main task. We converted spike times to firing rates as we have done in our prior studies (Chen et al. 2015). We then performed subsequent analyses on firing rates, as we describe next.

To measure stimulus-evoked visual responses, we aligned firing rates to grating onset. We grouped trials based on condition (e.g. 0.56 cpd gratings), and we ensured that there were no microsaccades occurring between -100 and 150 ms relative to stimulus onset. This was done in order to avoid other influences of microsaccades on SC evoked visual responses, which we have characterized elsewhere (Chen and Hafed 2017; Chen et al. 2015; Hafed and Krauzlis 2010). Specifically, when a stimulus onset occurs in the vicinity of a microsaccade onset, the subsequent stimulus-evoked response in SC neurons is modulated; our goal here was to characterize the unmodulated stimulus-evoked response and relate it to visual reafferent responses when a microsaccade jittered the image of a stably presented grating over the recorded RF.

When summarizing measurements across neurons, we measured peak firing rate in a given condition as the peak of the average firing rate of the condition across trials. That is, we grouped all trials of the same type, and we then averaged firing rates aligned on stimulus onset. We then defined a measurement interval 0-150 ms after stimulus onset during which we searched for a peak in the average firing rate curve. The peak value and peak time provided our estimate of peak firing rate of the neuron and visual response latency of the neuron.

When we visualized population firing rate curves across neurons in a given condition, we first normalized each neuron’s firing rate to a maximum of 1 before averaging the curves of the pooled neurons. Specifically, in a given condition (e.g. onset of a grating of 0.56 cpd spatial frequency), we pooled all trials and obtained an average firing rate curve. If the response to this condition was to be designated as the normalization reference, we then divided the firing rate curves of the same neuron in all individual trials (and all individual conditions) by the peak of this average firing rate curve. This way, all firing rates of the neuron across all trials and conditions were normalized to the firing rate of a given condition (e.g. 0.56 cpd). In different analyses, we either normalized by the peak visual evoked response to 0.56 cpd gratings (e.g. Fig. 5) or by the peak visual evoked response to the preferred spatial frequency of the neuron (e.g. Fig. 6). In this case, neuron “preference” was defined based on response strength (i.e. a preferred stimulus caused the strongest evoked response); in a lot of neurons, the preferred spatial frequency was indeed 0.56 cpd (Chen et al. 2018). It should also be noted here that in all analyses in which we presented normalized average firing rates, we also made sure to report raw activity measurements on a per individual neuron basis (e.g. Figs. 5, 6, 8, 9).

**Figure 3.**
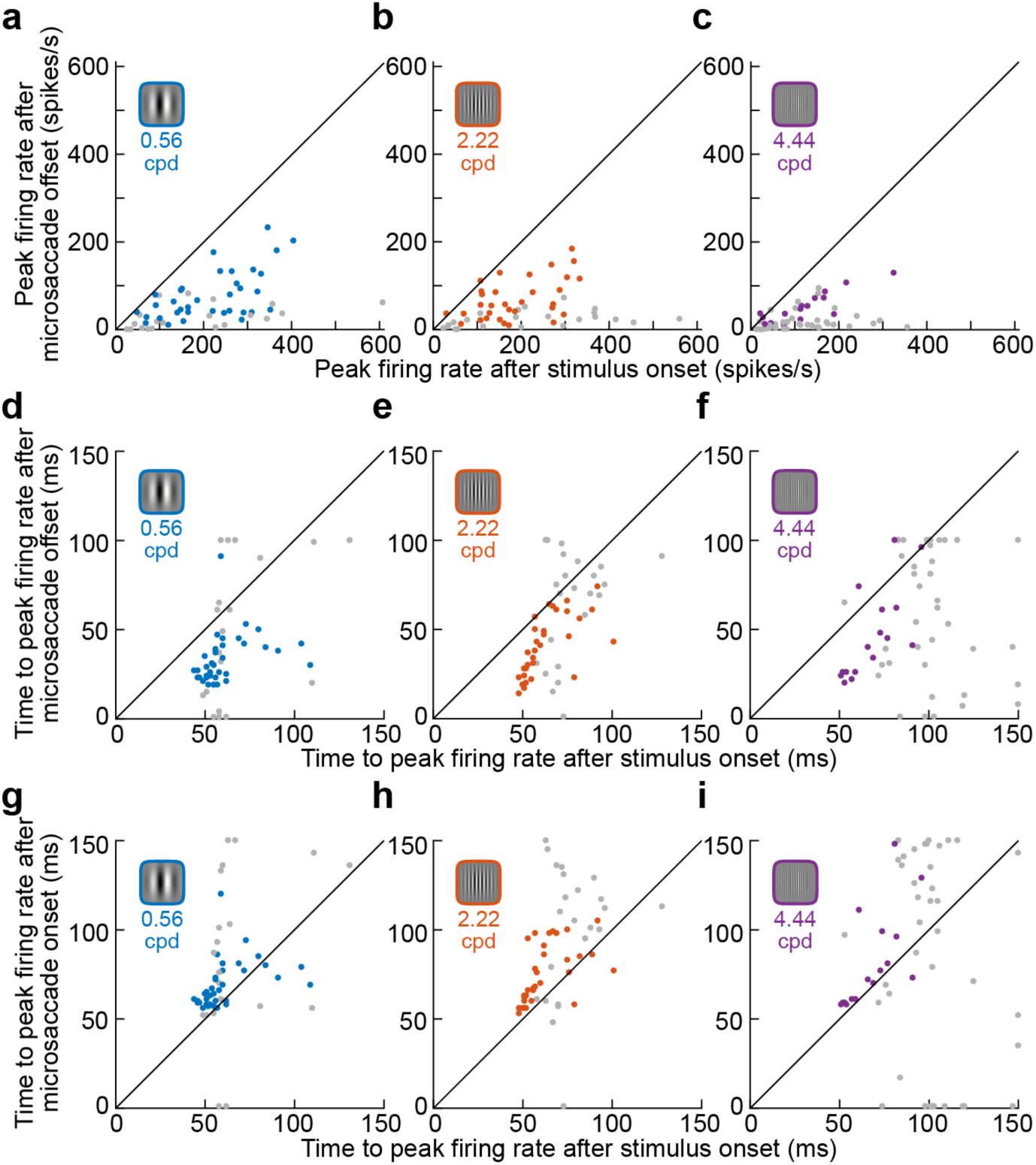
Properties of visual evoked and visual reafferent SC responses across the population. **(a)** For the lowest spatial frequency (0.56 cpd), we plotted peak firing rate after microsaccade end (the visual reafferent response) as a function of peak firing rate after stimulus onset (the visual evoked response). The gray symbols denote the neurons for which the visual reafferent response was not present (that is, statistically indistinguishable from baseline activity before microsaccades; Materials and Methods). **(b, c)** Same as **a** for the intermediate (**b**) and highest (**c**) spatial frequencies (2.22 cpd and 4.44 cpd, respectively). Visual reafferent responses were always weaker than visual evoked responses (p<0.0001, <0.0001, and <0.0001 across all neurons; t-test comparing reafferent to evoked responses for low, intermediate, and high spatial frequencies, respectively). **(d-f)** Same as **a**-**c** but now plotting the time to peak firing rate instead of the value of peak firing rate. For the visual reafferent responses, time to peak firing rate was measured from microsaccade end (p<0.0001, <0.0001, and <0.0001 across all neurons for low, intermediate, and high spatial frequencies, respectively). **(g-i)** Same as **d**-**f**, but measuring time to peak firing rate in the visual reafferent responses from microsaccade onset instead of microsaccade end (p=0.006, <0.0001, and 0.3 for low, intermediate, and high spatial frequencies, respectively).

**Figure 4.**
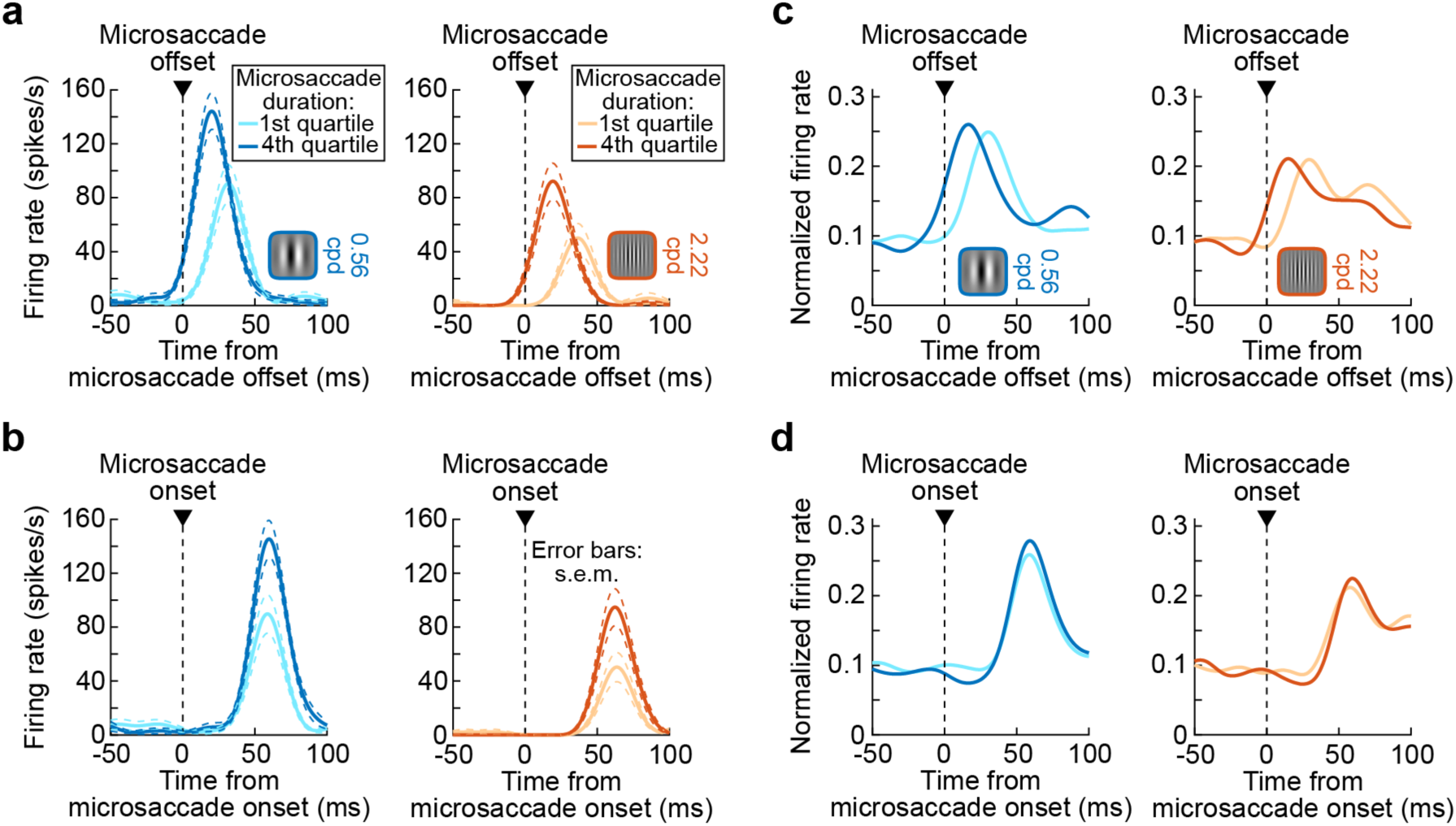
Alignment of visual reafferent responses to microsaccade onset rather than microsaccade end. **(a)** For the example neuron of Fig. 2, we binned microsaccades based on their duration. The reafferent response was clearly earlier (relative to microsaccade end) when microsaccades were longer in duration than when they were shorter in duration. Similar results were obtained for the highest spatial frequency (not shown), but the reafferent response was just weaker (Fig. 2). Error bars denote s.e.m. across microsaccades. **(b)** For the same neuron, aligning the same reafferent responses to microsaccade onset rather than microsaccade end revealed no delaying of responses for the short-duration microsaccades (even though the responses were significantly weaker than those for the long-duration microsaccades). Therefore, microsaccade-induced visual reafferent responses reflect a response to the entire movement event (starting with and including the initial motion blur period associated with eyeball rotation). **(c, d)** Similar results across our population. For each neuron, we binned microsaccade durations into 4 quartiles. We then averaged (across the population) responses for each quartile bin individually. Before averaging, we normalized each neuron’s response to its peak visual evoked response after the onset of the lowest spatial frequency (see Fig. 5). Consistent with **a**, **b**, reafferent responses were later relative to microsaccade end when the microsaccade was short in duration. When aligned on microsaccade onset, the same responses had the same latency (or were even slightly earlier) than the responses after long-duration microsaccades.

**Figure 5.**
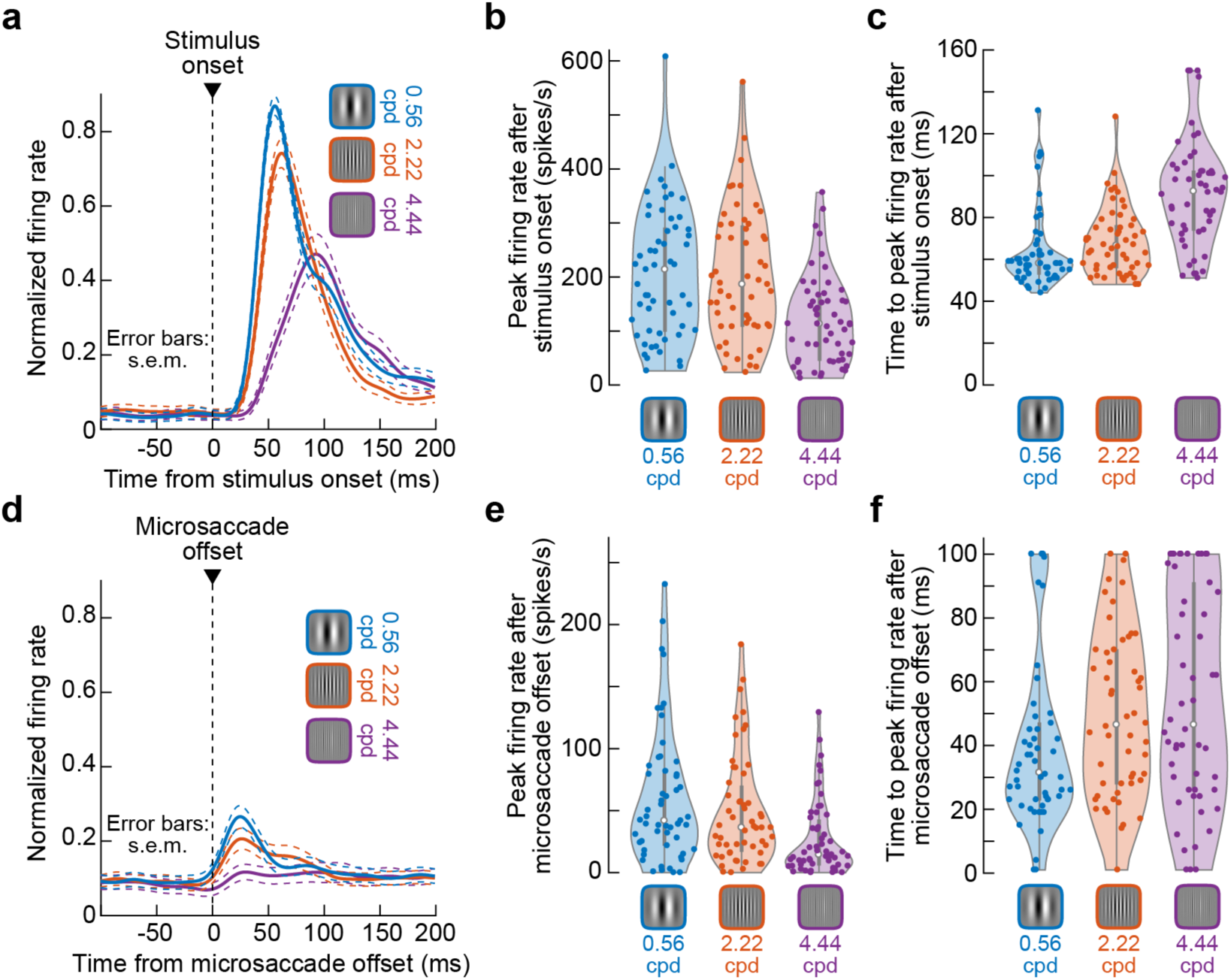
Comparison of response tuning to different spatial frequencies in either the visual evoked response or the visual reafferent response. **(a)** Firing rates of all neurons after stimulus onset. Each neuron’s firing rate curve was first normalized by the firing rate curve obtained for the lowest spatial frequency. Then, all neurons were combined (n=55 neurons). Error bars denote s.e.m. across neurons. As can be seen, the population preferred the lowest spatial frequency in terms of response magnitude and also response latency (shorter for lower spatial frequencies). **(b)** Individual neuron measurements of peak firing rate (unnormalized) after stimulus onset. The measurements (from the same neurons as in **a**) are separated based on stimulus spatial frequency. **(c)** For the same neurons, we now calculated time to peak firing rate, as a proxy for visual response latency. The visual evoked response was earliest for the lowest spatial frequency (Chen et al. 2018). **(d)** For the same neurons and the same normalization factor as in **a**, we plotted the visual reafferent response. Error bars denote s.e.m. across neurons. The visual reafferent response was weak, but still preferred the lowest spatial frequency. **(e, f)** Same as **b**, **c** but now for the visual reafferent response.

**Figure 6.**
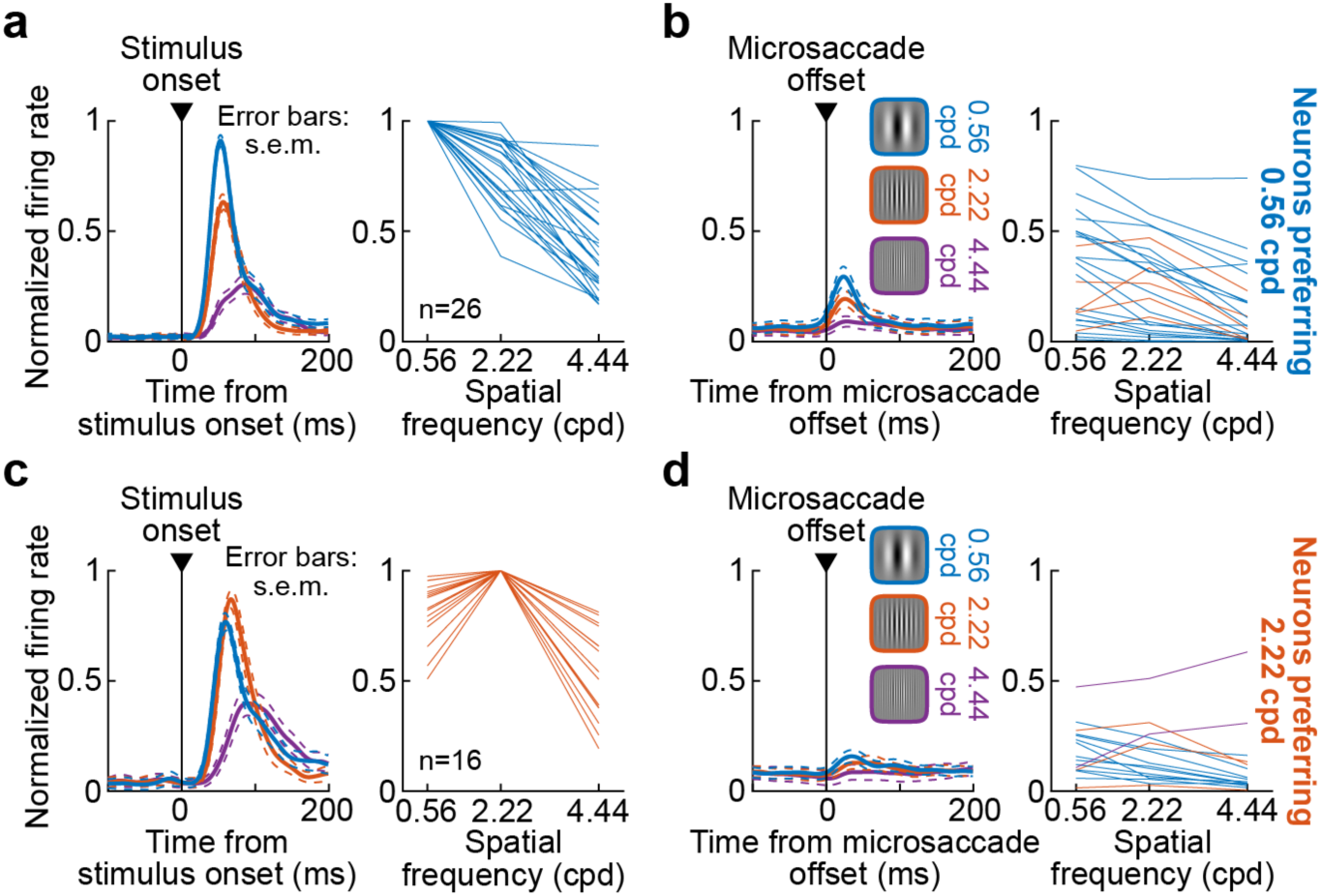
Differences in tuning between the visual evoked and visual reafferent responses of SC neurons. **(a)** The left panel shows normalized firing rate (similar to Fig. 5a) for all neurons preferring the lowest spatial frequency. Firing rates were normalized by the visual evoked response to the lowest spatial frequency (the preferred stimulus). Error bars denote s.e.m. across neurons. The right panel shows the individual neuron tuning curves. **(b)** For the same neurons, we plotted the visual reafferent response aligned on microsaccade end. The same normalization factor as in **a** was used. The right panel shows the tuning curves of the same neurons during the visual reafferent response. Most (21/26) neurons still preferred the lowest spatial frequency, but some preferred the intermediate spatial frequency (5/26; the differently colored tuning curves). **(c, d)** Same as **a**, **b** but now for the neurons preferring the intermediate spatial frequency. Firing rate curves were now normalized by the visual evoked response after presentation of the intermediate spatial frequency (the preferred stimulus). The tuning curves in **d** were different from those in **c**. Only 3/16 neurons still preferred the intermediate spatial frequency in the visual reafferent response; most (11/16) preferred the lowest spatial frequency instead.

**Figure 7.**
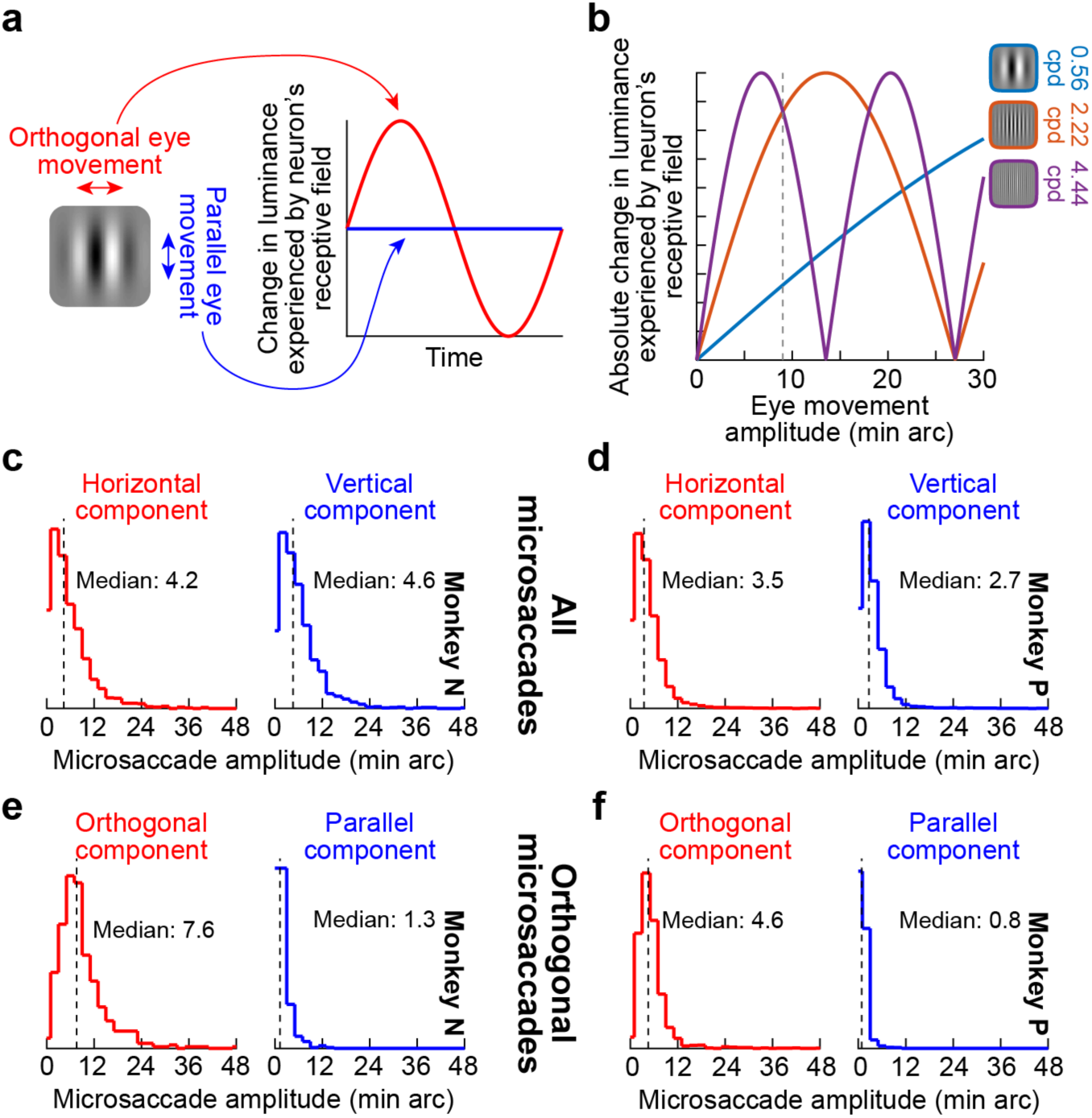
Potential impacts of microsaccade sizes and directions on properties of the visual reafferent response of SC neurons. **(a)** A movement orthogonal to grating orientation (horizontal in this case) is expected to cause luminance modulation in time as a result of eye-movement induced shift in retinotpic stimulus position; a parallel eye movement is expected to cause no modulation. Therefore, movement direction relative to stimulus orientation should modulate the visual reafferent response. **(b)** In addition, the amplitude of a movement in relation to the spatial frequency of the stimulus should cause predictable changes in luminance modulation. Here, assuming a purely orthogonal eye movement, the absolute value of amplitude modulation expected for a given spatial frequency is plotted as a function of eye movement amplitude. For example, for the lowest spatial frequency, a very large microsaccade is needed to cause a strong luminance modulation over a neuron’s RF (the dashed vertical line denotes 9 min arc). **(c, d)** Microsaccade amplitude probabilities like in Fig. 1e, but now plotted separately for the horizontal and vertical components of microsaccades. **(e, f)** Amplitude probabilities as in **c**, **d** but now for predominantly orthogonal (horizontal) microsaccades. These were movements with directions within +/- 22.5 deg from horizontal. The parallel component of these movements was, expectedly, much reduced. Predominantly parallel microsaccades had similar histograms (not shown), except that the small-amplitude distributions were now in the orthogonal, rather than parallel, component of the movements.

**Figure 8.**
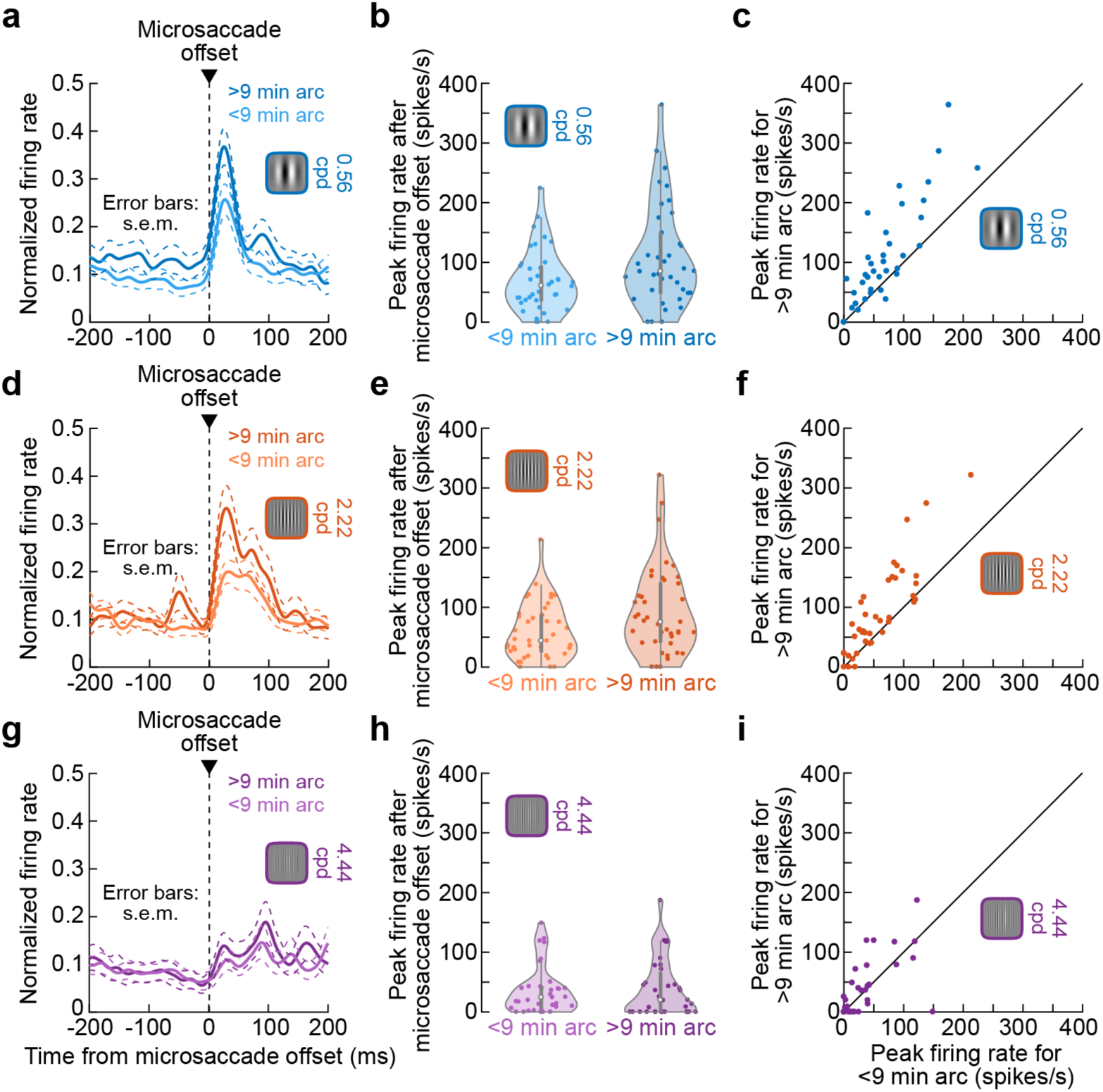
Dependence of the visual reafferent response on microsaccade size in relation to spatial pattern layout. **(a)** We plotted the average normalized visual reafferent response of neurons (normalized as in Fig. 5) when the RF stimulus had a spatial frequency of 0.56 cpd. The two curves indicate the reafferent response for microsaccades >9 min arc in amplitude or <9 min arc in amplitude, respectively. Error bars denote s.e.m. (across averaged neurons). **(b)** Individual neuron measurements for the same data as in **a**. The distribution of activity for larger microsaccades was higher than that for smaller microsaccades (p<0.0001; t-test). **(c)** The same data but now plotted as paired measurements, showing enhanced an visual reafferent reafferent response for the larger microsaccades. **(d-f)** Similar results for the intermediate spatial frequency (p<0.0001; t-test). **(g-i)** For the highest spatial frequency, there was no clear difference in the strength of the visual reafferent response as a function of larger or smaller microsaccades (p=0.61; t-test).

**Figure 9.**
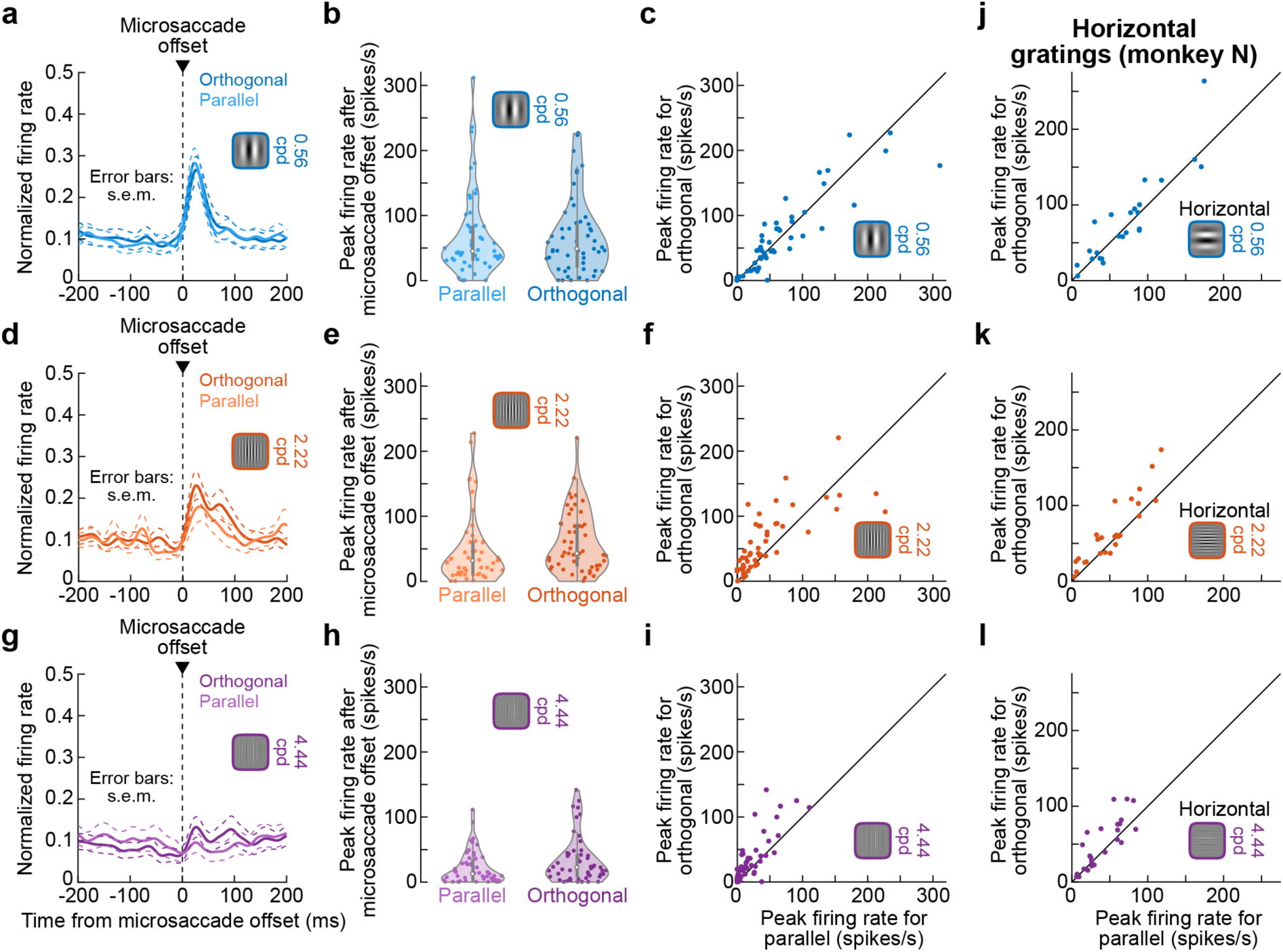
Dependence of the visual reafferent response on microsaccade direction relative to the orientation of luminance modulation patterns in a visual stimulus. **(a-i)** These panels are formatted identically to Fig. 8, except that the comparison is now between microsaccades that were predominantly orthogonal to the grating in the RF or microsaccades that were predominantly parallel (i.e. predominantly horizontal microsaccades versus predominantly vertical microsaccades). Orthogonal microsaccades were associated with a stronger visual reafferent response than parallel microsaccades, except for the lowest spatial frequency (for which our observed microsaccade sizes were too small to result in appreciable differences in luminance modulations; Fig. 7b). P-values for statistical comparisons for each spatial frequency are reported in the text. **(j-l)** To confirm that the results were related to the relative relationships between microsaccade direction and stimulus orientation (as opposed to, say, absolute microsaccade direction irrespective of the stimulus orientation), we repeated the same analyses on recordings from additional experiments performed with horizontal gratings in monkey N. Now, “orthogonal” means predominantly vertical microsaccades. The same conclusions as in **a**-**i** were reached: firing rates were higher for orthogonal than parallel microsaccades, except for the lowest spatial frequency (p=0.22, 0.0036, and 0.0184, respectively, for increasing spatial frequencies). The panels are otherwise formatted identically to **c**, **f**, **i**.

To statistically determine whether a neuron had a significant visual response for a given spatial frequency compared to baseline activity, we defined two measurement intervals. The first was 0-150 ms before stimulus onset and served as the baseline firing rate of the neuron. The second was 0-150 ms after stimulus onset and served as the visual evoked epoch. Across repetitions of a given condition, if the average firing rate in the second interval was statistically higher (at a p-value of <0.01) than the average firing rate in the first interval (Wilcoxon rank sum test if numbers of repetitions were <16; t-test otherwise), then the neuron possessed a statistically significant visual response.

For some analyses, we were also interested in whether individual neurons possessed statistically significant tuning to an individual spatial frequency (from among the three that we sampled in our experiments). To do that, we defined an interval of 0-150 ms after stimulus onset. For each neuron and spatial frequency, we estimated the peak firing rate per trial. We then performed a one-way ANOVA with the main factor being spatial frequency. If the ANOVA showed a significant main effect of spatial frequency (at a p-value of <0.05), we also performed a post-hoc t-test (at a p-value of <0.01). This allowed us to group the neurons into four groups: preferring the lowest spatial frequency (0.56 cpd); preferring the intermediate spatial frequency (2.22 cpd); preferring the highest spatial frequency (4.44 cpd); or not preferring any of the three presented spatial frequencies (no significant main effect in the one-way ANOVA).

To measure visual reafferent responses, we used the same procedures above, but we aligned firing rates to microsaccade end (that is, new fixation onset). When we normalized firing rate curves (see above), we always normalized to visual evoked, not visual reafferent, responses. This allowed us to quantitatively compare the strength of visual reafferent SC responses relative to stimulus-driven visual evoked responses. Also, when statistically assessing whether there was a significant visual reafferent response, we used the same concept of two measurement intervals as described above. However, in this case, the “baseline” interval was 50-150 ms before microsaccade onset, and the “response” interval was 0-100 ms after microsaccade end. The latter interval was early (compared to visual evoked measurement intervals above) because the visual reafferent responses had shorter latencies from microsaccade end (see Results).

In all analyses, we only analyzed visual reafferent responses for microsaccades that occurred when the grating was already stably present on the display (that is, already present inside the neuron’s RF; see Results). Specifically, we only considered microsaccades occurring from 300 ms after grating onset until 150 ms before trial end. The former time value was to ensure that the visual evoked response after sudden stimulus onset had subsided; the latter time value was to ensure that we had enough trial repetitions across all collected data.

To further understand and support our observations from spiking activity as assessed with firing rate measurements, we also analyzed local field potentials (LFP’s) as a highly sensitive population measure of SC visual tuning properties around the recording electrodes (Chen and Hafed 2017; 2018; Chen et al. 2018; Hafed and Chen 2016; Ikeda et al. 2015); LFP’s were additionally shown to be useful in analyses from other brain areas beyond the SC (e.g. Neupane et al. 2016). We used similar procedures to those described earlier (Chen and Hafed 2017; 2018; Chen et al. 2018; Hafed and Chen 2016). Briefly, we started with the wide-band signals digitized at 40 KHz by our data acquisition system, which also initially applied a hardware filter passing the range of 0.7 Hz to 6 KHz frequencies. We then removed electrical power line noise artifacts using IIR notch filters at 50, 100, and 150 Hz. Finally, we applied a zero-phase-lag low pass filter with cutoff frequency of 300 Hz, and we down sampled the resulting data to 1 KHz. This gave us continuous signals of LFP modulations analogous to continuous firing rate curves obtained from individual spike times. For each recording site, we averaged the individual-trial LFP curves and aligned them to either stimulus onset or microsaccade onset/end.

To ensure robustness of our interpretations, we performed within-neuron analyses, and we report summaries from each neuron individually. We also enforced minimum numbers of trials for analyses as described above. When separating neurons according to their preferred spatial frequencies (e.g. Fig. 6), we only showed spatial frequencies for which there were enough neurons preferring them to allow reasonable interpretations of the results (i.e. with enough sampled neurons).

Given the strong congruence of the visual evoked responses (to stimulus onsets) with our recent observations on SC spatial frequency tuning properties (Chen et al. 2018), we are confident that our numbers of neurons are sufficient to represent the overall SC population and summarize our observations on visual reafferent responses. Moreover, the use of two monkeys is viewed as a minimal compromise between using animals in research, on the one hand, and ensuring replicability of results across larger populations of research subjects, on the other.

We report measures of variance for all analyses and figures, and we also show individual neuron results for all analyses. Statistical tests are reported at relevant portions of Results, and in separate analyses (not shown), we also further investigated all of our statistical tests using alternative methods involving randomization tests.

When reporting p-values, we used the following conventions for purposes of clarity: if a p-value was less than 0.0001, we reported it simply as p<0.0001; if a p-value was higher, we reported its exact value.

### Data availability

All data presented in this paper are stored in institute computers and are available upon reasonable request.

## Results

We explored the visual consequences of fixational microsaccades on neural activity in the rhesus macaque SC. We did so by presenting a Gabor grating of a given orientation and spatial frequency at a visual location corresponding to the visual RF location of a recorded neuron (Fig. 1a). The grating was stationary and persisted on the display for up to approximately 3000 ms, but it was continuously moved relative to the recorded neuron’s retinotopic RF by occasional microsaccades. Across sessions, we recorded from primarily parafoveal neurons in both the right and left SC (average preferred eccentricity: 3.5 deg +/- 0.39 deg s.e.m.; n=55 neurons; vertical grating experiments), and the grating radii were tailored in each experiment according to the recorded RF size and location. We explored the visual reafferent response after a microsaccade jittered the grating over the recorded neuron’s RF. In all cases, we analyzed the activity of visually-responsive SC neurons, whether they also had a saccade-related motor burst or not, but we ensured that none of our neurons had a motor burst for microsaccades in any direction (Hafed et al. 2009; Willeke et al. 2019). Otherwise, it would have been difficult to differentiate between whether neural activity after microsaccades was related to visual reafference or to the microsaccade-related discharge itself. Figure 1b demonstrates the lack of peri-microsaccadic neural discharge that we aimed for in our population of neurons when no visual stimulus was present inside the recorded neuron’s RF.

We also ensured that microsaccade directions across our experiments were independent of RF location (Fig. 1c). This was to be expected because the stimuli were irrelevant to the rewarded behavior (fixation), and because our prior work demonstrated that microsaccades in these types of tasks correct for fixational eye position errors (foveal motor errors) even in the presence of peripheral cues (Tian et al. 2018; 2016). We also found a similar result to Fig. 1c earlier (Chen et al. 2015). Because we fixed grating orientation across RF locations, this independence of microsaccade directions from RF locations (Fig. 1c) also meant that our microsaccade directions were additionally independent of grating orientations.

### Microsaccades are associated with a visual reafferent response in SC

We first compared visual reafferent responses after microsaccades to responses when the grating first appeared inside a neuron’s RF. As shown in Fig. 1a, each monkey fixated a central fixation spot (Materials and Methods), and we presented a vertical grating filling the RF; RF size was initially assessed with small spot stimuli in initial RF mapping tasks (Materials and Methods). The example trial shown in Fig. 1d demonstrates how stimulus onset elicited a short-latency visual response, and how the neuron continued to fire occasionally afterwards, and particularly after microsaccades. We characterized the post-stimulus and post-microsaccadic neural responses. The great majority of microsaccades that we analyzed had a radial amplitude of <0.2 deg (<12 min arc; Fig. 1e), and they were associated with peak velocities significantly less than approximately 70 deg/s (medians: 10 deg/s and 23 deg/s for monkeys P and N, respectively).

For each presented spatial frequency, we plotted the visual evoked response of the same example neuron of Fig. 1d (Fig. 2a), as well as the visual reafferent response after microsaccades (Fig. 2b); Figure 2c shows the same visual reafferent response as in Fig. 2b, but now aligned to microsaccade onset rather than microsaccade end (an indication of microsaccade duration variability is portrayed in the figure by the gray vertical line and its surrounding shaded region). For the visual reafferent response (Fig. 2b, c), we only analyzed microsaccades for which the grating was already displayed to the monkey. That is, we picked a sustained time interval starting 300 ms after grating onset and ending 150 ms before trial end, and we only analyzed microsaccades during this interval. This interval started after the initial stimulus-driven visual evoked response of the neuron had subsided, such that the visual reafferent response was analyzed when there was already a stable stimulus inside the RF (the microsaccades were small enough to not move the presented stimulus outside the RF when they occurred).

As can be seen from the visual evoked response (Fig. 2a), this example neuron preferred the lowest presented spatial frequency (0.56 cpd), emitting the strongest peak firing rate. Moreover, the neuron’s visual response latency also increased with increasing spatial frequency, consistent with recent observations in the SC (Chen et al. 2018). In the visual reafferent response (Fig. 2b, c), the neuron’s activity was weaker, but it maintained the preference for the lowest spatial frequency. Specifically, the peak firing rate after microsaccades for the lowest spatial frequency (0.56 cpd) was 104 spikes/s when aligned to microsaccade end; this was 38% of the peak firing rate for the same spatial frequency after stimulus onset. For the intermediate spatial frequency (2.22 cpd), the peak firing rate of the neuron after microsaccades was 62 spikes/s (31% of the stimulus-evoked response), and the peak firing rate for the highest spatial frequency (4.44 cpd) was 12 spikes/s (18% of the stimulus-evoked response). Thus, the firing rate after microsaccades was not simply a scaled-down version (by a constant factor across spatial frequencies) of the stimulus-evoked response. Also note that due to variability of microsaccade durations, aligning on microsaccade end (new fixation onset) rather than microsaccade onset resulted in marginally weaker visual reafferent responses in the former alignment relative to the latter (compare panels in Fig. 2b, c). For example, for the lowest spatial frequency (0.56 cpd), the peak visual reafferent response was 104 spikes/s when aligned on microsaccade end and 112 spikes/s when aligned on microsaccade onset. We will briefly return to this observation shortly, when describing response latencies in more detail (e.g. Fig. 4).

Across the population, we measured the visual evoked response after stimulus onset as the peak firing rate emitted by a neuron during an interval 0-150 ms after stimulus onset (Materials and Methods). We also measured the visual reafferent response as the peak firing rate during an interval 0-100 ms after microsaccade end; the intervals were chosen based on the timings of bursts after stimulus onsets or microsaccades observed across our population (Materials and Methods). Almost every single neuron that we recorded from showed a weaker visual reafferent response than a visual evoked response (Fig. 3a-c). In fact, the visual reafferent response was often seemingly absent (especially for the grating with the highest spatial frequency), even when a neuron responded after stimulus onset.

To statistically test whether the visual reafferent response was present or not in a given neuron, we compared, for each neuron, average firing rate in the interval 0-100 ms after microsaccade end (the visual reafference epoch) to average firing rate in the interval 50-150 ms before microsaccade onset (the baseline epoch). If the post-microsaccade firing rate across repetitions was higher than the pre-microsaccade firing rate at a p-value of <0.01 (assessed using a t-test or ranksum test; see Materials and Methods), we deemed the neuron to possess a significant visual reafferent response. Otherwise, the neuron lacked a significant visual reafferent response. The neurons colored in gray in Fig. 3a-c were the neurons without any significant visual reafferent response. For the lowest spatial frequency (0.56 cpd; Fig. 3a), these non-responding neurons comprised 36% of our recorded population (20/55); for the intermediate spatial frequency (2.22 cpd; Fig. 3b), this fraction was 42% (23/55); and, finally, for the highest spatial frequency (4.44 cpd; Fig. 3c), 71% (39/55) of our recorded neurons showed no visual reafferent response after microsaccades. Note that for the visual evoked response after stimulus onset, all neurons that we analyzed showed a response for all spatial frequencies. Thus, the reafferent response was sometimes weak enough to simply disappear completely. However, this might be expected: the highest spatial frequency (which showed the least likelihood of significant visual reafferent response) was anyway associated with the weakest visual evoked response overall (compare the x-axis ranges of Fig. 3a-c) (Chen et al. 2018).

The results above suggest, so far, that the SC exhibits visual reafferent responses after microsaccades, similar to other early visual areas (Bosman et al. 2009; Herrington et al. 2009; Kagan et al. 2008; Leopold and Logothetis 1998; Martinez-Conde et al. 2002; 2000; Snodderly et al. 2001). The strength of these responses can be variable, and even absent, in some cases. We next describe the latencies of these responses before returning to the potential reasons for such variability, and complete absence in some cases.

### Visual reafferent responses after microsaccades are earlier than visual evoked responses after stimulus onset

Since a new fixation begins after the end of the prior eye movement, visual reafferent responses are frequently aligned in analyses on movement end (that is, new fixation onset; e.g. Kagan et al. 2008). With such a convention, visual reafferent responses after microsaccades occurred in our data set significantly earlier than visual evoked responses after stimulus onset. This is already evident in the example neuron of Fig. 2. As can be seen, the time to peak firing rate after stimulus onset in this neuron was 44 ms, 52 ms, and 66 ms for the low, intermediate, and high spatial frequencies, respectively. On the other hand, after microsaccade end, these times were 27 ms, 28 ms, and 40 ms instead. Across the population, the visual reafferent response (when it did occur) had a latency (to peak firing rate) of 33 ms +/- 2.5 ms s.e.m. from microsaccade end for the lowest spatial frequency, 40 ms +/- 3 ms s.e.m. for the intermediate spatial frequency, and 46 ms +/- 6 ms s.e.m. for the highest spatial frequency (versus 61 ms +/- 2 ms s.e.m., 62 ms +/- 3 ms s.e.m., and 68 ms +/- 6 ms s.e.m. for the stimulus-driven visual evoked response, respectively). Thus, the post-microsaccadic visual reafferent response was ∼24 ms earlier than the visual evoked response (Fig. 3d-f). These results are consistent with other studies for large saccades in other areas (e.g. Ibbotson et al. 2008; Kagan et al. 2008; Price et al. 2005; Rajkai et al. 2008; White et al. 2017b).

Note that the latencies in Fig. 2 and Fig. 3d-f after microsaccades might appear too fast given visual conduction delays in the early visual system. However, the stimulus jittering that happens on the retina because of a microsaccade (or saccade) actually starts with the movement onset itself. That is, neurons do not merely respond to the stimulus at the start of the new fixation, but they respond to the entire spatiotemporal pattern of stimulus movement caused by the eye movement. Therefore, it is expected that the visual reafferent response is affected by the entire event of a microsaccade, including the initial motion component, and not just reacting to the new stable image of the grating after the end of the eye movement. Consistent with this, when we computed the timing of the visual reafferent response now relative to microsaccade onset rather than microsaccade end, we found that it had similar or longer latency than the original stimulus-driven visual evoked response (Fig. 3g-i). This delaying (Fig. 3g-i) and weakening (Fig. 3a-c) of the visual reafferent response (when compared to the stimulus-driven visual evoked response) is expected due to the motion blur associated with the microsaccadic movements themselves.

To further confirm that the SC reafferent response was likely a response to both the motion blur caused by microsaccades and the new stable image inside the RF, we hypothesized that the reafferent response latency should depend on microsaccade duration when aligned on microsaccade end, but be independent of microsaccade duration when aligned on microsaccade onset. That is, if the response is due to the entire event of motion blur plus stable image, then the response should come earlier relative to microsaccade end if the motion blur duration is slightly elongated (due to a longer-duration microsaccade). This is indeed what we found. In Fig. 4a, b, we analyzed the activity of the same example neuron as that shown in Fig. 2 after binning microsaccades based on their duration; specifically, we binned microsaccades into four duration quartiles, and we plotted reafferent responses as a function of quartile, either aligned on microsaccade end (Fig. 4a) or microsaccade onset (Fig. 4b; the figure only shows the first and fourth quartiles for purposes of clarity, but the other quartiles gave expected intermediate results). The reafferent response was clearly earlier relative to microsaccade end for longer-duration microsaccades (Fig. 4a). In this example neuron, the response was also stronger because longer-duration microsaccades tend to also be larger in size (see Fig. 8 below for why microsaccade size matters).

Across the population, we found similar results. For each neuron, we binned microsaccade durations into 4 quartiles. We then plotted normalized firing rate (normalized to the peak response of the stimulus-evoked response to the lowest spatial frequency; Materials and Methods and Fig. 5) after microsaccades for each group, and we averaged the population neural responses. There was a clear rank ordering of reafferent response latency as a function of microsaccade duration, but only when aligned on microsaccade end (Fig. 4c, d; once again, the figure shows the first and last quartile results only for clarity, but the other quartiles gave expected intermediate results). Note that the effect is not explained by the possibility that stronger responses merely come earlier than weak responses, because alignment on microsaccade onset still revealed stronger responses for longer-duration microsaccades but without a response latency difference (Fig. 4b, d). These observations are also consistent with our earlier observations in Fig. 2 that the visual reafferent response was stronger when aligned on microsaccade onset rather than microsaccade end.

### Visual reafferent responses after microsaccades do not always reflect the visual tuning of SC neurons assessed with the evoked response

The visual consequence of any given eye movement should depend on the size and direction of the movement relative to the stimulus pattern (that is, how the spatial luminance modulation in the stimulus gets translated over a given RF). This means that the visual reafferent response might not always be a replica (albeit weakened; Fig. 3) of the visual evoked response. We investigated this issue, which is under-represented in the neurophysiology literature of post-saccadic reafference, in our recorded SC neurons.

We first asked whether response strength and response latency obeyed known observations about SC visual responses to Gabor gratings (Chen et al. 2018). In the visual evoked response, we found that most of our neurons (47%) preferred the lowest spatial frequency in terms of firing rate magnitude, consistent with (Chen et al. 2018). To demonstrate this, we normalized each neuron’s visual evoked response to the peak firing rate observed after the onset of the grating with the lowest spatial frequency (0.56 cpd). We then divided all firing rate measurements across trials, and across spatial frequencies, by this measurement. We also used the same normalization factor for the visual reafferent response. We then pooled all neurons’ normalized firing rates in Fig. 5a for the visual evoked response and in Fig. 5d for the visual reafferent response (for simplicity, we only show the reafferent response aligned on microsaccade end, rather than also aligned on microsaccade onset, in this figure). In terms of response magnitude, tuning was generally maintained in the visual reafferent response. That is, the visual reafferent response was strongest for the lowest spatial frequency and weakest for the highest spatial frequency. This idea is also shown in Fig. 5b, e, in which individual neuron distributions are presented under the different stimulus presentation conditions of low, intermediate, and high spatial frequencies; there was a significant main effect of spatial frequency (one-way ANOVA) in the visual evoked response (F(2,162)=10.89, p<0.0001) and also the visual reafferent response (F(2,162)=7.89, p=0.0005).

Interestingly, the rank ordering of response latencies with increasing spatial frequency, which was observed earlier (Chen et al. 2018), was also present in the visual reafferent response. Specifically, in Fig. 5c, we plotted the time to peak firing rate in the visual evoked response of neurons, and confirmed the earlier observations (Chen et al. 2018). In Fig. 5f, we plotted the time to peak firing rate of the visual reafferent response after microsaccade end. The range of measurements was naturally broader and more variable than for the visual evoked response (Fig. 5c), but this is expected given the variable durations and speeds of individual microsaccades (introducing variability in both the amount and duration of the motion blur caused by eye movements before a new “fixation” was eventually established; Fig. 4).

Nonetheless, there was a clear trend for increasing response latency with increasing spatial frequency, as in the visual evoked response (also seen in the example neuron of Fig. 2): there was a significant main effect of spatial frequency (one-way ANOVA) in the visual evoked response (F(2,162)=30.27, p<0.0001) and also the visual reafferent response (F(2,162)=4.54, p=0.0121).

These results indicate that even though microsaccades are very small (Fig. 1e) and relatively slow eye movement events (significantly less than 70 deg/s peak velocity), they still refresh SC activity to represent a stimulus that was moved over the neuronal RF’s by them. Such refreshing is mild (Fig. 3), but still present (Fig. 5), and it is generally faithful to the visual evoked tuning of neurons.

Having said the above, at the individual neuron level, we did observe cases of the tuning of a given neuron apparently changing in the visual reafferent response when compared to the visual evoked response. For example, we grouped the neurons that preferred the lowest (26 neurons) or intermediate (16 neurons) spatial frequency. We then repeated the same analyses of Fig. 5a, d but this time normalizing firing rates to the peak firing rate of the preferred spatial frequency (rather than to the lowest spatial frequency as in Fig. 5a, d). Such new normalization strategy was now necessary to assess whether tuning (i.e. preferred spatial frequency) was altered. Neurons preferring the lowest spatial frequency in the visual evoked response still primarily preferred the lowest spatial frequency in the visual reafferent response (Fig. 6a, b). Specifically, the left panel of Fig. 6a shows an analysis similar to that of Fig. 5a. In the right panel of the same figure, we plotted the tuning curves of individual neurons for the three different spatial frequencies. All neurons in this group preferred the lowest spatial frequency in the stimulus-driven visual evoked response (as per the design of the analysis). Now, for the visual reafferent response after microsaccades (Fig. 6b), the tuning curves of the same neurons largely still preferred the lowest spatial frequency stimulus (with a milder reafferent response strength; Fig. 3). However, some neurons actually responded in a stronger fashion for the intermediate spatial frequency (2.22 cpd) rather than the lowest spatial frequency (5/26; 19%). This effect was even more striking when we repeated the same tuning curve analysis for neurons preferring the intermediate spatial frequency in the visual evoked response; these neurons often preferred the lowest spatial frequency in the visual reafferent response (Fig. 6c, d); that is, the visual reafferent response was strongest for the lowest rather than the intermediate spatial frequency (11/16 neurons; 69%). A similar effect occurred for the neurons preferring the highest spatial frequency, although there were too few neurons in this condition (especially with a significant visual reafferent response); we therefore do not show the results for the neurons preferring the highest spatial frequency.

These results suggest that when the visual reafferent response is weak (that is, for the intermediate and highest spatial frequencies), then it may be most sensitive to interactions between the microsaccadic eye movement’s vector and the stimulus properties, resulting in an apparent change in visual feature tuning of the neuron during the visual reafferent response. We directly demonstrate the evidence supporting this hypothesis in the next section; in other words, there is visual coding of neural patterns caused by the specific microsaccadic eye movement vectors.

### Visual reafferent responses after microsaccades depend on the temporal image luminance modulations over RF’s caused by the shifting eye positions

To clarify the potential importance of luminance modulations caused by a given eye movement for the results of Fig. 5 above, we further investigated the relationships between microsaccade properties and stimulus properties. Specifically, in the experiments described so far, the grating was vertical (Materials and Methods). This means that a purely horizontal microsaccade would cause luminance modulation over the RF as the eye movement displaces the retinotopic image of the grating horizontally (orthogonal to the grating), whereas a purely vertical microsaccade would not cause such luminance modulation (Fig. 7a). Moreover, depending on the spatial frequency of the grating, the size of the shift caused by a purely horizontal microsaccade should influence the absolute amplitude luminance modulation experienced at a given retinotopic RF location (Fig. 7b). For example, for a horizontal microsaccade and a vertical grating of 0.56 cpd, the largest luminance modulation over an RF expected by the eye movement would occur for a large microsaccade of approximately 0.9 deg amplitude (54 min arc) (Fig. 7b). Therefore, both microsaccade size and microsaccade direction should influence the strength of the visual reafferent response in SC neurons, resulting in an apparent change in tuning in some cases, as we saw in Fig. 6. We tested for this idea by isolating microsaccades of specific sizes and directions.

To do so, we first inspected the constraints on expected luminance modulations that were imposed on us by our observed microsaccade amplitudes (Fig. 1e). Both monkeys had microsaccades of different directions, having different sizes of horizontal or vertical microsaccade components (Fig. 7c, d). This allowed us to develop specific expectations (Fig. 7b) on the impacts of, say, microsaccade size or microsaccade direction on the visual reafferent response to a given spatial frequency (e.g. 0.56 cpd). We first investigated the effects of microsaccade size, and we then moved on to next investigate the impacts of microsaccade orientation relative to grating orientation. For the latter, we picked microsaccades with angles that were within +/- 22.5 deg from either horizontal or vertical, to isolate movements that were either primarily orthogonal or primarily parallel to the vertical grating, respectively. For example, in Fig. 7e, f, we measured the distributions of predominantly orthogonal microsaccades (very small “parallel” amplitude component in the figure) in each monkey, which we could use in conjunction with Fig. 7b to predict potential effect sizes for different stimulus spatial frequencies.

In terms of microsaccade size, we picked all horizontal microsaccades (within +/- 22.5 deg from horizontal in direction), and we binned their amplitudes as being either smaller or larger than 0.15 deg (9 min arc; this binning was, once again, constrained by our observed microsaccade amplitude distributions in Figs. 1e, 7c-f). Given the predicted luminance modulation caused by a given movement size (Fig. 7b), one should expect that the larger microsaccades would be associated with larger visual reafferent responses, but only for the lowest and intermediate spatial frequencies. Specifically, between the amplitudes of 9 min arc and ∼20-22 min arc (the upper limit of the great majority of our microsaccades; Fig. 1e), the expected luminance modulations caused by the eye movements are higher for both low and intermediate spatial frequencies than the expected luminance modulations caused by smaller microsaccades with amplitudes <9 min arc (Fig. 7b). For the highest spatial frequency, the same ranges of microsaccade amplitudes that were available to us (Figs. 1e, 7c-f) would not be expected to reveal any systematic effect. This was indeed the case. In Fig. 8a, d, g, we plotted the visual reafferent response for “small” (<9 min arc) and “large” (>9 min arc) horizontal microsaccades, across the different stimulus spatial frequencies. The violin plots (Fig. 8b, e, h) show raw measurements of the visual reafferent response (peak firing rate of a neuron in the interval 0-100 ms from microsaccade end), and the firing rate curves (Fig. 8a, d, g) show mean curves across the population of neurons (after each neuron’s firing rate curve was first normalized by the peak visual response after stimulus onset for the lowest spatial frequency; as in Fig. 5 above). Error bars denote s.e.m. across neurons. As can be seen, the visual reafferent response was stronger for larger microsaccades (within the range of microsaccade amplitudes that we were able to analyze), but not for the highest spatial frequency. We confirmed this statistically in Fig. 8c, f, i by plotting the measurements in Fig. 8b, e, h as paired observations (reafferent response after large microsaccades versus reafferent response after small microsaccades). The clearest effects were present for the low and intermediate spatial frequencies but not for the highest spatial frequency (p<0.0001, p<0.0001, and p=0.61, respectively; t-test). Therefore, consistent with the predictions of Fig. 7b, microsaccade size had a clear influence on post-microsaccadic visual reafferent responses in the SC, and in a manner that demonstrated dependence on temporal luminance modulations over the RF caused by the eye movements.

In terms of microsaccade directions, we expected that movements orthogonal to the gratings (that is, predominantly horizontal movements) should have the strongest visual reafferent response since they cause the strongest luminance modulations of the gratings over the recorded RF (Fig. 7a). We therefore separated microsaccades according to whether they were predominantly horizontal or predominantly vertical (Fig. 9a-i). In each case, and as stated above, we defined predominantly horizontal or predominantly vertical as all movements with directions within +/- 22.5 deg from the designated direction. The strongest visual reafferent response occurred for the horizontal microsaccades (labeled orthogonal in the figure since the gratings used were vertical), but (unlike with microsaccade size effects in Fig. 8) this effect was now strongest for the intermediate and high spatial frequencies and not for the lowest spatial frequency. Specifically, in both the average firing rate curves (Fig. 9a, d, f) as well as the individual neuron distributions (Fig. 9b, e, h), we could observe stronger visual reafferent responses for orthogonal compared to parallel microsaccades in the intermediate and high spatial frequencies; also see the paired measurements of the individual neuron distributions shown in Fig. 9c, f, i (p=0.0114, 0.0002 for the intermediate and high spatial frequencies, respectively; t-test). However, there was no difference in the visual reafferent response when the microsaccades jittered the image of a low spatial frequency grating either horizontally or vertically (orthogonally or in a parallel fashion; p=0.63; t-test). Given the overall sizes of our microsaccades (Figs. 1e, 7c-f), this was expected because the lowest spatial frequency would have required substantially larger eye movements to cause strong luminance modulations (Fig. 7b).

We also confirmed that the results of Figs. 8, 9a-i remained valid even when we restricted the analyses to only neurons preferring the lowest spatial frequency in their visual evoked response (e.g. Fig. 6) or only neurons preferring the middle spatial frequency (data not shown). That is, the modulatory influence of microsaccade amplitude and direction was present and the same regardless of the original stimulus-evoked feature preference of a neuron. More importantly, we also wondered whether the results of Fig. 9a-i were not related to movement direction relative to stimulus orientation per se (i.e. not related to “orthogonal” versus “parallel”) but to absolute movement direction (i.e. horizontal versus vertical microsaccades). We therefore conducted additional experiments testing horizontal as opposed to vertical gratings in one monkey (monkey N). With such gratings, we expected the biggest reafferent responses to occur after predominantly vertical microsaccades rather than after predominantly horizontal microsaccades, and we also expected that the effect of microsaccade direction should depend on spatial frequency (Fig. 9a-i). This is indeed what we found, and with similar statistical conclusions as a function of spatial frequency (Fig. 9j-l).

Therefore, to summarize our results so far, we found that a significant component of the effects of microsaccades on vision is to modulate how SC visual neural activity represents stimuli that are already stably present in the environment (Figs. 1-5). The interaction between the visual transient caused by the eye movement (that is, the retinotopic motion caused by the movement) and the movement vector itself relative to the underlying stimulus pattern will vary how SC neural activity is modulated (Figs. 7-9), and therefore how image representations are ultimately decoded downstream depending on the needed behavioral readout of SC activity. This can give the potential appearance of changes in visual tuning of SC neurons during the visual reafferent response interval (e.g. Fig. 6).

We finally aimed to further support the above interpretations by analyzing LFP signals around our recording electrodes. We previously saw that LFP measurements in the SC provide a highly sensitive measure of population activity (Chen and Hafed 2017; 2018; Chen et al. 2018; Hafed and Chen 2016). We therefore analyzed visual evoked LFP deflections across our population, as well as microsaccade-aligned LFP deflections (Materials and Methods). We first replicated our recent observations that the visual evoked LFP response preferred low spatial frequencies (Chen et al. 2018). This replication is shown in Fig. 10a (in a similar format to that used in Chen et al. 2018); stronger LFP deflections (more negative) occurred for the lowest spatial frequencies. We then aligned the LFP signals on microsaccade end (Fig. 10b) or microsaccade onset (Fig. 10c), similar to how we aligned firing rates earlier (e.g. Fig. 2). We found that when a microsaccade happened with a stable visual stimulus near the aggregate population RF location around the recorded sites, there was a clear visual reafferent LFP response. Interestingly, even though the reafferent response was weaker than the visual evoked response (compare the y-axis ranges in Fig. 10a and Fig. 10b, c), there was clear feature tuning of the reafferent response: the visual reafferent LFP response was strongest (most negative) for the lowest spatial frequency, and progressively weakened with increasing spatial frequency (Fig. 10b, c), just like the visual evoked LFP response (Fig. 10a). Moreover, we found that the post-microsaccadic LFP response was a genuine reafferent response because the LFP signal aligned on microsaccades when there was no visual stimulus on the display at all had a much weaker deflection (inset in Fig. 10c; also see Chen and Hafed 2017). Therefore, our LFP analyses supported the notion of visual feature tuning of reafferent neural responses in the SC after microsaccades.

**Figure 10.**
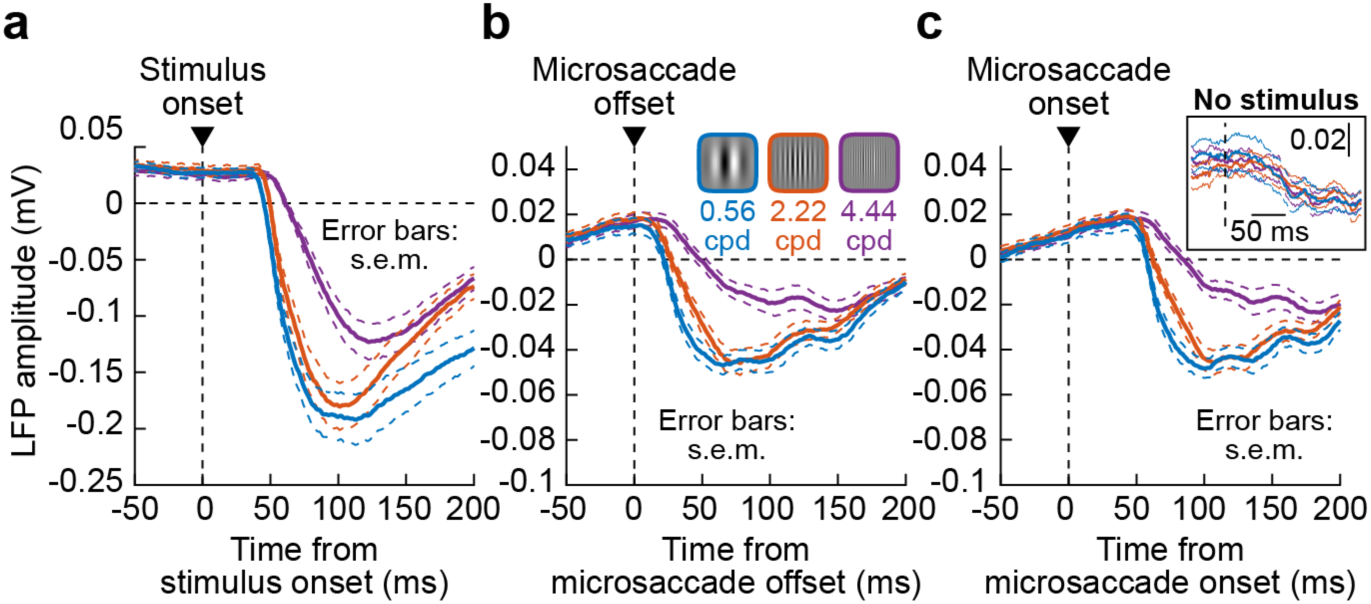
A visual reafferent response in LFP signals in the primate SC. **(a)** We plotted the average stimulus-driven LFP deflection across all of our recorded SC sites (error bars denote s.e.m. across sites). Each colored curve shows the LFP response after the onset of a given spatial frequency (color coded as in the legend in **b**) (n=55 sessions; vertical grating experiments). The LFP signal reflected a general population preference for low spatial frequencies in the visual evoked response (Chen et al. 2018). **(b)** For the same recording sites, we aligned the LFP signal to microsaccade end when a grating of a given spatial frequency (different colors) was in the aggregate RF location represented by the recorded SC site for a given session. Consistent with the firing rate data (e.g. Figs. 2-5), there was a clear visual reafferent LFP response, which also reflected the feature tuning of the visual evoked LFP response in **a**. That is, there was a stronger response (stronger negativity) after microsaccades for low spatial frequencies. Note that the visual reafferent LFP response, despite showing similar feature tuning to **a**, was weaker in strength (compare y-axes in **a**, **b**). **(c)** Same data as in **b**, but now aligned on microsaccade onset. Similar observations were made. The inset shows that when microsaccades occurred in the absence of a visual stimulus on the display, the microsaccade-aligned LFP response was much weaker (less negative) than when a stimulus was present. This confirms that the results in **b**, **c** were genuine reflections of SC reafferent responses. The different colors in the inset denote that stimulus-absent microsaccades were from trials in which a stimulus of a given spatial frequency (color coded) would later appear.

More importantly, when considering the impacts of microsaccade size as a function of spatial frequency, exactly like we did in our firing rate analyses (Fig. 8), we found consistent results in the visual reafferent LFP response (Fig. 11a-c): there was a stronger reafferent LFP response for larger rather than smaller microsaccades (that were orthogonal to the gratings), but only when low and intermediate spatial frequencies were being jittered on the retina by the eye movements. Microsaccade size (for our observed range: Figs. 1e, 7c-f) did not matter when high spatial frequencies were stably present in the environment. As stated above, these results are consistent with expectations of temporal luminance modulations caused by microsaccades, for the given range of movement amplitudes that was present in our data (Fig. 7a, b).

**Figure 11.**
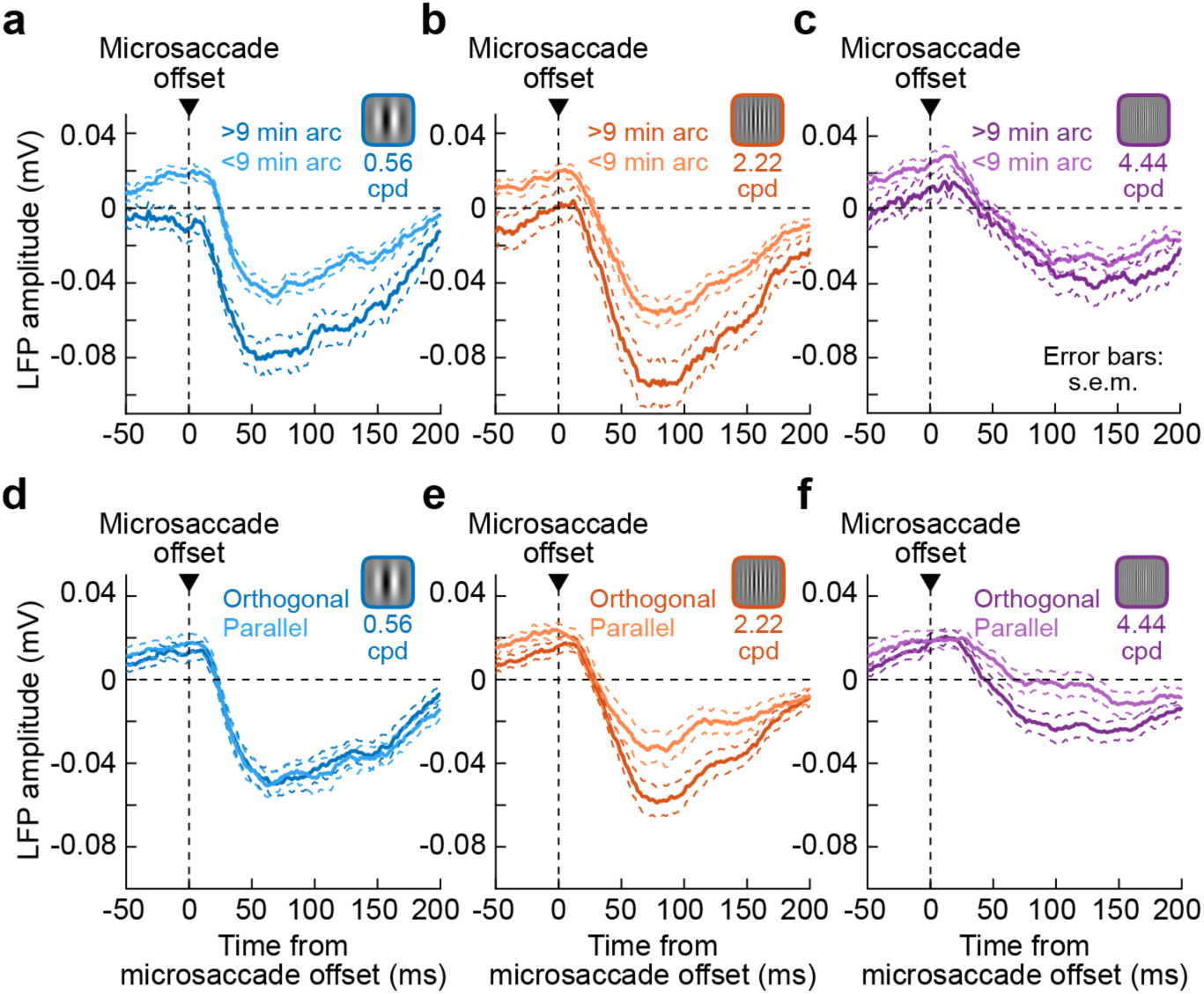
Dependence of the visual reafferent LFP response on microsaccade size and direction relative to spatial pattern layout. **(a-c)** Similar analyses to those in Fig. 8, but this time on microsaccade-aligned LFP signals. For each spatial frequency, we compared the visual reafferent LFP response when the microsaccade that jittered the displayed stimulus was >9 min arc in amplitude or <9 min arc in amplitude. There was a stronger visual reafferent LFP response for the larger microsaccades, except for the highest spatial frequency (consistent with Figs. 7b, 8). Error bars denote s.e.m. across sessions. **(d-f)** Similar analyses to those in Fig. 9a-i, but this time on microsaccade-aligned LFP signals. The comparison is now between microsaccades orthogonal or parallel to the displayed stimulus, as opposed to larger or smaller as in **a**-**c**. Consistent with Figs. 7b, 9, there was a clearly stronger visual reafferent LFP response for the intermediate and high spatial frequencies, but not for the low spatial frequency.

Similarly, when we separated microsaccades based on their directions (orthogonal and parallel) relative to the gratings, the same predictions (Fig. 7a, b) suggested stronger reafferent responses for orthogonal rather than parallel gratings, but only for intermediate and high spatial frequencies. This was, again, clearly the case in our visual reafferent LFP responses (Fig. 11d-f), just like in Fig. 9 for spiking activity. Therefore, both our firing rate and LFP analyses demonstrated that the SC continuously updates its representation of stably present stimuli in the visual environment, and particularly with periodic post-microsaccadic reafferent responses that critically depend on interactions between the movement vector (dictating temporal modulations of luminance over an RF), on the one hand, and the spatial layout of the stimuli, on the other.

## Discussion

We investigated the properties of the visual reafferent response in SC neurons after microsaccades jittered stable stimuli inside the neurons’ RF’s. We found that the SC exhibits a visual reafferent response that is clearly visible in firing rates and LFP signals. The visual reafferent response is significantly weaker than the stimulus-driven visual evoked response, and sometimes also absent. Moreover, the visual reafferent response exhibits feature tuning properties generally similar to those of the visual evoked response. However, at the individual neuron level, some neurons may appear to alter their feature tuning preferences in the visual reafferent response when compared to the visual evoked response. Investigating the detailed interactions between microsaccade sizes/directions and image patterns led us to conclude that such alteration of feature tuning likely reflects the impacts of the temporal modulations of luminance caused over an RF by a microsaccade. Such modulations depend not only on the spatial frequency of the stimulus itself, but also on the eye movement vector (size/direction).

Our results indicate that the SC provides a continuous representation of a visual scene, with periodic “updating” through visual reafferent responses even with tiny microsaccades. In recent work, we investigated foveal visual neurons in the SC, and we found that these neurons reflect how slow fixational drift eye movements can bring the fixated spot in and out of the neurons’ RF’s (Chen et al. 2019). This causes variability in the neurons’ firing activity during fixation. The current results go beyond such previous work by showing that even parafoveal and peripheral SC neurons continue to represent visual scenes as they are jittered on the retina by microsaccades. This is interesting because it demonstrates that the visual properties of the SC that seem to be very relevant for orienting behavior, such as RF size and visual sensitivity (Chen et al. 2018; Hafed 2018; Hafed and Chen 2016), also extend to visual representations when stimuli are stable in the environment and moved retinotopically by eye movements. It would be interesting in future studies to explore how the SC’s representation of stable environments can influence subsequent visuo-motor behavior, exactly through its visual reafferent responses. For example, do enhanced reafferent responses after microsaccades facilitate subsequent saccades in the same direction as the extra-foveal bursting neurons?

One interesting consequence of results like those shown in Fig. 6, with apparent changes in feature tuning of some neurons after microsaccades, is that there may be inherent ambiguities in individual neurons’ response strengths as a consequence of eye movements. While it may be conceivable that individual neurons can change their feature tuning curves across eye movements, similar in principle to how spatial RF’s can be significantly altered around the time of saccades (Duhamel et al. 1992; Walker et al. 1995; Zirnsak et al. 2014), we think that a more likely source of the alterations that we observed in our experiments (Fig. 6) was the expected temporal luminance modulation over the recorded RF caused by eye movements (Figs. 7-9, 11). Specifically, we found that there were predictable dependencies of the visual reafferent response strength on both the stimulus spatial profiles as well as the eye movement vectors. This functionally means that eye movements contribute to visual coding by enhancing some feature properties (Rucci et al. 2018; Rucci et al. 2007; Rucci and Victor 2015), and in our case, this happened even for microsaccades. For example, edges in an image orthogonal to the microsaccades would be enhanced significantly more than edges parallel to the microsaccades (Fig. 9, 11d-f). More interestingly, such visual coding also happens within the SC, a structure that has been primarily thought of for many years as being more of a motor control structure rather than a visual structure.

Related to the above, another interesting consequence of our results is that the post-microsaccadic SC visual reafferent response may impose on peripheral SC neurons a temporal tag of updated visual representations that is somewhat independent of position coding (because of the small sizes of the eye movements). Specifically, in natural viewing, very small saccades may be frequently required to scan features of far objects (e.g. features of a mountain peak viewed with the naked eye on a hiking adventure, or features of the far road ahead when driving at high speed on a highway). Because the RF’s of peripheral SC neurons, particularly in the lower visual field, are relatively big (Hafed and Chen 2016), such tiny saccades (Fig. 1e) are too small to move the images being jittered on the retina by the eye movements outside or away from the neurons’ RF’s. Nonetheless, according to our results, the temporal modulations caused by the small retinal jitter would still cause a reafferent response in the peripheral neurons. Therefore, peripheral edges or other visual features may be enhanced periodically by small saccades in natural viewing, even when these small saccades may be used for high acuity foveal tasks (like inspecting far objects). This periodic “updating” maintains the peripheral image representations that may be needed to re-orient gaze with larger eye movements to specific peripheral features of interest. It also means that the SC visual reafferent response that we studied after microsaccades here is not restricted to being a highly constrained laboratory phenomenon with enforced fixation like in our tests. Rather, it can still happen in many natural scenarios in which very small saccades would be frequently employed anyway.

Combined with results from other visual brain areas (Bosman et al. 2009; Herrington et al. 2009; Kagan et al. 2008; Leopold and Logothetis 1998; Martinez-Conde et al. 2002; 2000; Snodderly et al. 2001), the above temporal tagging idea that we alluded to (i.e. the periodic updating of image processed representations by microsaccadic or larger saccadic eye movements) may in reality be an almost global phenomenon across the whole brain. Indeed, several brain areas that may be concerned with other sensory (e.g. auditory) or motor (e.g. arm movement) modalities are also sensitive to visual stimulation. As a result, and because vision comes through a mobile eye, these same brain areas may be sensitive to eye-movement-induced visual stimulation due to retinal image movements (i.e. potential visual reafferent responses). This means that visual processing is almost never immune from the impacts of individual saccadic or microsaccadic eye movements. In fact, eye movements cause phase-resetting of visual and oculomotor systems (Bellet et al. 2017; Gaarder et al. 1966; Hafed and Ignashchenkova 2013), and there are very long-lasting impacts of such resetting on perception (Bellet et al. 2017). All of this leads to an urgent conclusion that vision, by virtue of its continually mobile sensor, cannot and should not be studied in isolation of orienting behaviors, in general, and eye movements, in particular.

Finally, a component of eye movements that still occurs in the absence of saccades and microsaccades is that of slow ocular drifts. Such fixational drifts are related to microsaccades and saccades, because drift properties are clearly related to, and potentially affected by, the occurrence of the latter more rapid eye movements (e.g. (Chen and Hafed 2013). Moreover, drifts continue to occur in between microsaccades, even when strict gaze fixation is required. This means that the same idea of temporal luminance modulations over an RF that we described in this study would still be valid for SC neurons with slow ocular drifts. However, given the much smaller sizes of drift eye movements (∼ 1 min arc scale) and their much lower speeds (< 0.5 deg/s or less) when compared to even microsaccades, the amount of displacements and potential motion blur associated with drift eye movements would be expected to be much smaller than even those associated with the very small movements that we encountered in our experiments (Figs. 1e, 7). It remains to be seen whether SC visual responses would be sensitive to such minute, almost indiscernible, image modulations associated with slow ocular drift eye movements. In our future experiments, this will be explicitly tested, and particularly with causal manipulations of real-time retinal image stabilization (and careful eye movement recording) in order to allow us to establish even more strong evidence that SC neurons provide a continuous, real-time representation of our visual environment that is reformatted by active oculomotor behavior during fixation.

## Acknowledgements

We were funded by the Deutsche Forschungsgemeinschaft (DFG, German Research Foundation; projektnummer 276693517—SFB 1233). We were also funded by the Werner Reichardt Centre for Integrative Neuroscience (DFG Excellence Cluster: EXC 307) and the Hertie Institute for Clinical Brain Research.

